# Improved cortical boundary registration for locally distorted fMRI scans

**DOI:** 10.1101/248120

**Authors:** Tim van Mourik, Peter J Koopmans, David G Norris

## Abstract

With continuing advances in MRI techniques and the emergence of higher static field strengths, submillimetre spatial resolution is now possible in human functional imaging experiments. This has opened up the way for more specific types of analysis, for example investigation of the cortical layers of the brain. With this increased specificity, it is important to correct for the geometrical distortions that are inherent to echo planar imaging (EPI). Inconveniently, higher field strength also increases these distortions. The resulting displacements can easily amount to several millimetres and as such pose a serious problem for laminar analysis. We here present a method, Recursive Boundary Registration (RBR), that corrects distortions between an anatomical and an EPI volume. By recursively applying Boundary Based Registration (BBR) on progressively smaller subregions of the brain we generate an accurate whole-brain registration, based on the grey-white matter contrast. Explicit care is taken that the deformation does not break the topology of the cortical surface, which is an important requirement for several of the most common subsequent steps in laminar analysis. We show that RBR obtains submillimetre accuracy with respect to a manually distorted gold standard, and apply it to a set of human in vivo scans to show a clear increase in spacial specificity. RBR further automates the process of non-linear distortion correction. This is an important step towards routine human laminar fMRI. We provide the code for the RBR algorithm, as well as a variety of functions to better investigate registration performance in a public GitHub repository, https://github.com/TimVanMourik/OpenFmriAnalysis, under the GPL 3.0 license.

## Introduction

Investigation of the BOLD response with functional MRI at the level of the cortical layers has become increasingly popular over the the last decade (Dumoulin et al., 2017; Trampel et al., 2017). Activation levels differ at the laminar scale (Koopmans et al., 2011) and they can vary depending on the performed task (Muckli et al., 2015; Kok et al., 2016). Laminar signals have the potential to reveal information about the underlying neuronal processes within a cortical region, as the signal from different layers may be associated with feed forward or feedback signals (Felleman and Van Essen, 1991; Self et al., 2017). However, layer specific analysis comes with great methodological challenges.

The thickness of the cerebral cortex varies between 1 and 4.5 millimetres (Zilles, 1990; Fischl and Dale, 2000). Identifying individual layers therefore ideally requires sub-millimetre resolution, at the cost of signal to noise ratio (SNR) per voxel. On top of this, a functional experiment ideally requires a Repetition Time (TR) on the order of several seconds. Layer specific investigations are hence best conducted at higher field (≥7 Tesla) for improved SNR, but high field strength also have some disadvantages (Poser and Setsompop, 2017). The inhomogeneities of the static magnetic field *B*_0_ can cause non-linear distortions when a fast acquisition scheme like Echo Planar Imaging (EPI) is used (Mansfield, 1977). Distortions primarily present themselves in the phase-encoding direction as a function of the bandwidth per pixel and static field strength (Schmitt et al., 1998). As layer specific analysis requires high spatial precision, non-linear distortions are particularly problematic.
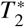
-weighted images usually have insufficient contrast to segment the cortical grey matter, so instead one needs to identify the cortical boundaries from a different scan. This is typically a high-contrast *T*_1_-weighted anatomical scan that can be segmented with tools such as FreeSurfer (Dale et al., 1999), CBS Tools (Bazin et al., 2014), or BrainVoyager (Goebel, 2012). However, as the anatomical scan is undistorted, it may not sufficiently overlap with the functional scan.

Several potential solutions have been proposed for providing accurate alignment of functional images with an anatomical image. Early papers circumvent the problem by segmenting only a small piece of straight cortex (Ress et al., 2007; Koopmans et al., 2010). It is also possible to resort to different acquisition schemes like FLASH, which do not suffer significant distortions, but at a heavy cost in temporal resolution. Koopmans et al. combine this with a vertex based realignment procedure based on the Stripe of Gennari (Koopmans et al., 2011). This approach, however, is highly specific to parts of the primary visual cortex that show a myelinated band in the middle of the cortex (Stripe of Gennari), and does not generalise to the rest of the brain. Yet another alternative is to acquire an anatomical image with the same EPI readout and field of view as the functional image, such that the two volumes are similarly distorted (Kashyap et al., 2017). However, this (often unjustly) assumes that field inhomogeneities, and with it the distortions, do not change between acquisitions. Additionally, if the acquisition only covers a small part of the brain, cortical reconstruction algorithms may easily fail, as they are often based on whole-brain templates.

Ideally, there would be an accurate cross-contrast 
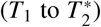
 registration algorithm, but this is a notoriously hard problem. The combination of the warping of images and the unknown relation between contrasts creates a vast parameter space that is difficult to solve in a coregistration procedure. While algorithms like AFNI’s 3dQWarp (Cox, 1996) technically support cross-modal cost functions, the documentation acknowledges that such usage is rather experimental and in our experience indeed does not reach submillimetre accuracy. In general, no algorithm currently exists that corrects non-linear distortions in high resolution low-contrast images up to the laminar specific level, and works either on partial or whole-brain images.

One way to search through relevant information in the image domain is to try and detect the grey-white matter boundary and match this between volumes. This is using *geometric* information on top of *volumetric* information and forms the basis of Boundary Based Registration (BBR) (Greve and Fischl, 2009). A three-dimensional cortical reconstruction of the grey-white matter boundary is created on a high contrast anatomical image and serves as a basis for the coregistration. While a functional image is too low in contrast for generating a cortical reconstruction, it can be used in the registration procedure. The contrast is sufficient to optimise the average contrast across the boundary in order to achieve a better realignment. This was proposed for linear registrations and proven to be an exceedingly robust method. We here extend BBR to work recursively and effectively produce a cross-contrast non-linear registration. Our aim was to provide accurate submillimetre registration for whole-brain or partial brain images. The algorithms can be performed on the the functional data, without additional acquisition of additional scans, and without reinterpolation as a result of a non-linear warping. Importantly, the procedure produces a smooth deformation field that does not alter the topological properties of the mesh, such that the resulting surfaces can naturally be used in subsequent automatic layering and further layer specific analyes (Waehnert et al., 2014; Leprince et al., 2015).

## Methods

Boundary Based Registration (BBR) is based on a cortical construction of the boundary between the white matter and grey matter, and the grey matter and the CSF (pial surface) (Greve and Fischl, 2009). This surface is generated on an anatomical scan, as the contrast of the functional scan may not be good enough for segmentation. In order to register the two volumes, the constructed surface is moved to the functional image. The average contrast across the grey-white matter boundary is computed and used as a cost function to optimise the transformation parameters of the registration. Versions of BBR are implemented in FreeSurfer (`bbregister`) (Dale et al., 1999) and FSL (`flirt -cost bbr`) (Jenkinson and Smith, 2001). This method is powerful enough to be able to register the volumes, based on only a small part of the brain (Greve and Fischl, 2009), suggesting that local application can also be successfully employed. Especially for high resolution (laminar) fMRI, the anatomical and functional volumes contain detailed information about the gyrification that can be used for registration. We hence propose to extend BBR with a hierarchical strategy (Collins et al., 1995). By recursively applying BBR at diminishing scales as a series of linear transformation, the volumes are effectively non-linearly registered.

The way in which the mesh is divided is based on a three-dimensional cuboid lattice consisting of the set of neigbourhoods 
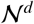
 at various depths *d*, decreasing in size. At each depth, 
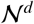
 is divided into eight equally sized cuboids (two in each dimension) 
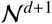
 until a user specified threshold size is reached. All vertices that make up the brain mesh are divided over the elements of 
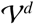
 the set of vertices within 
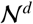
 The number of vertices in each neighbourhood may well vary between regions.

Registration is performed iteratively, from the largest scale (*d* = 0) to the smallest (*d* = *d_max_*). For each element 
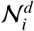
 a registration is computed by means of a boundary based registration algorithm (Greve and Fischl, 2009) applied to 
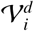
 In short, this is an edge detection algorithm that maximises the average contrast across the white matter surface. Contrast is defined as the (optionally weighted) gradient of samples on either side of vertices within the mesh. For the optimisation procedure we use MATLAB’s gradient descent method `fminsearch` (default parameters) to optimise registration parameters as a function of contrast. The same sampling method and cost function were used as proposed by Greve & Fischl (Greve and Fischl, 2009).

Applying the computed transformations directly to 
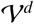
 could easily ‘break the mesh’ at the edges of neighbouring regions as their continuity is not guaranteed. This is a serious problem, as subsequent steps in laminar analysis (like the level set methods (Sethian, 1999)) require the mesh to be a topological sphere: a non-intersection closed surface without holes. In order to ensure continuity we use a control point based strategy (Collins et al., 1995): let the edges and corners of 
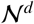
 define a deformable lattice. For each computed transformation in 
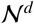
 a resulting displacement vector is assigned to all of its corner points. After all transformations at a depth level are computed, the median is taken of all displacement vectors for each control point, thus representing a resultant vector based on adjacent neighbourhoods. In order to further increase robustness (but at the cost of specificity), the displacement vectors may subsequently be adjusted based on their direct neighbours:

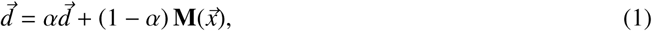

where 
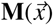
 denotes the mean displacement of the neighbourhood. Collins et al. (Collins et al., 1995) experimentally found *α* = 0.5 yields an acceptable balance between local matching and global smoothness. We recommend higher values for *α*, as our primary goal in laminar analysis is the local matching. We suggest additional ways of increasing robustness in the next section.

As all control points within 
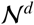
 now have displacement vectors associated with them, 
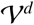
 an be displaced proportionally to the distance to their closest control points. A way of describing this is by defining transformation matrix **T**, such that it satisfies:

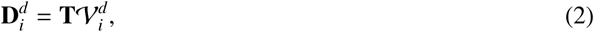

where the set of displacement vectors is denoted by **D***^d^*, for which each *i*’th element contains 8 vectors, one for each corner of the cube. This is a typical linear regression equation and could hence trivially be solved for **T**. However, a least squares solution is required as the system is overdetermined with eight corner points and only a [4*X*4] transformation matrix. This necessarily requires an approximation of the deformation vectors that may result in discontinuities with respect to adjacent neighbourhoods. A preferred method is to divide the cube into six tetrahedra by means of Delaunay triangulation (Delaunay, 1934) and compute a separate matrix for each of them. For each tetrahedron, equation 2 represents a determined system, for which the solution for adjacent tetrahedra is guaranteed to be continuous. Effectively, this division increases the degrees of freedom of the deformation field to satisfy our requirement of continuity. Having found the transformations, they can readily be applied to adjust the position of 
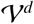
.

This procedure is repeated for all depths levels *d*. Whenever the number of vertices in 
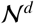

is smaller than a user specified minimum, the transformation for that region is set to the identity matrix.

## Robustness

With the exponential increase in number of neighbourhoods as a function of depth level, the number of registrations easily reaches into the hundreds. The high number of Degrees of Freedom (DoF) of the algorithm therefore inescapably increases the probability of misregistrations. The algorithm ensures robustness and continuity in several ways, by computing displacement vectors as a weighted average of all surrounding neighbourhoods, and by an (optional) additional smoothing of displacement vectors based on adjacent control points, as mentioned above. Additionally, we compute a transformation within a neighbourhood not only for the entire neighbourhood, but also for six subregions. By splitting 
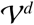
 separately into two in the *x*, *y*, and *z* dimensions, six cuboids are created for which the registration is repeated. This procedure is illustrated in Fig. 1. The resulting displacement vectors are added to the respective list for each control point.

**Figure 1:**
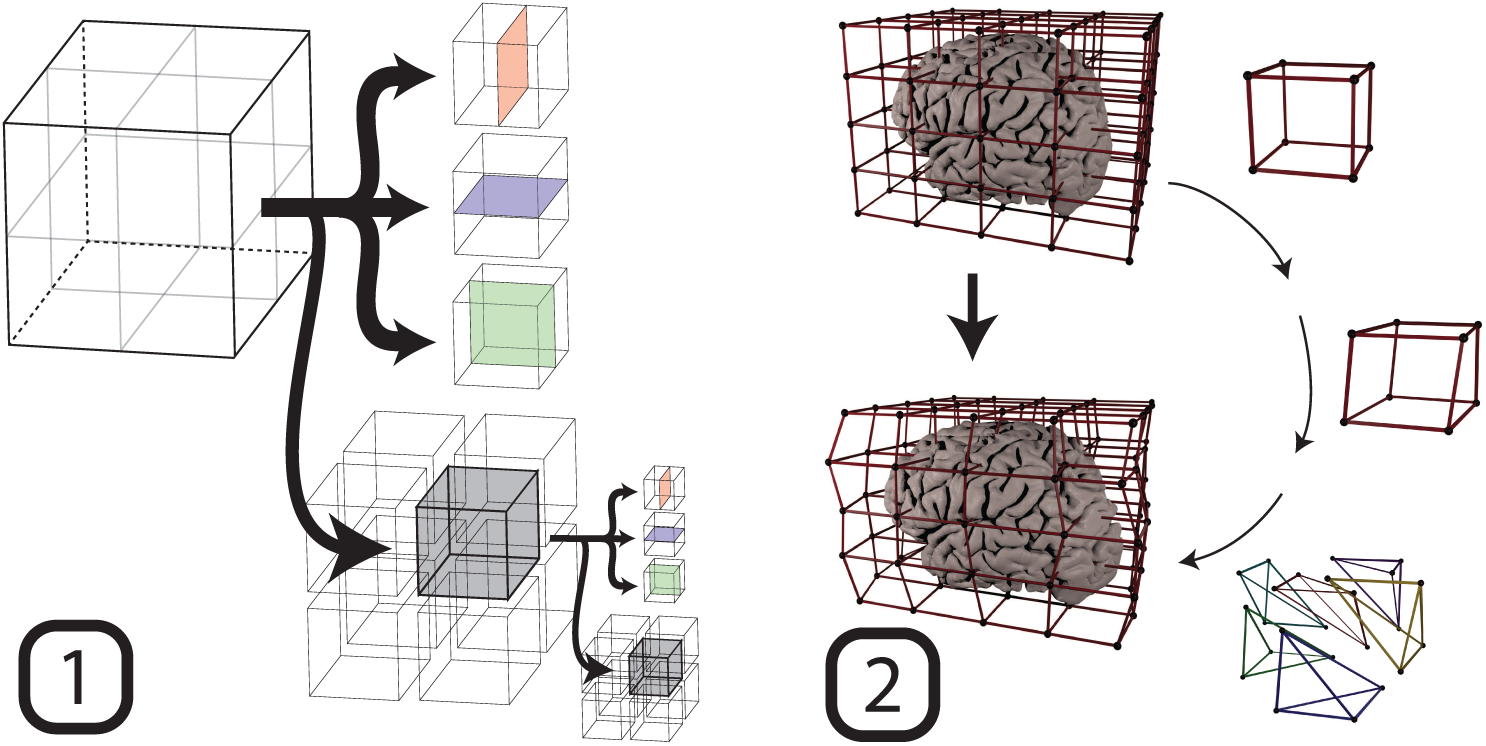
A schematic of the workflow. First, the volume is recursively broken up into parts, for which registrations are computed by means of the edge detecting boundary based registration method. Based on the transformations found in this step, the second step is the updating of a cuboid lattice. The control points within the lattice are updated and applied to the boundaries. Specifically, this was performed based on a tetrahedral division of the cube. For each tetrahedron, a displacement field was computed and applied to the vertices within the tetrahedron.

## Parameters

The algorithm can look for any combination of translation, rotation and scaling in the *x*, *y*, and *z* dimensions. It will divide the volume until it reaches a user defined minimum size or number of vertices, and a transformation will be computed. We here focus on registration in the phase enconding direction only, i.e. translation and scaling in the *y*-direction, as this is the most common type of distortion when an Echo Planar Imaging sequence is used (Mansfield, 1977). We set the minimum size of a neighbourhood to 4 voxels and the minimum number of vertices to 100. The smoothing factor with respect to adjacent vertices in the lattice from Eq.1 was set to *α* = 0.9. Experimentally, we have found that this still yields robust results while still being highly weighted towards specificity. In contrast, Collins et al. (Collins et al., 1995) describe an *α*-level of *α* = 0.5, which is likely to do with the fact that their purpose of template matching and segmentation prioritises robustness over specificity. While the original implementation of BBR recommends using subsets of vertices for parts of the algorithm (Greve and Fischl, 2009), we here use all vertices at all stages. This may be redundant for registrations at the large scale, but as it is a small addition in computation time, and because errors in early iterations may propagate further downwards, we chose to incorporate all vertices. Finding the best selection of parameters may still require some iterations for a given data set. It is, however, a substantial improvement with respect to the arduous job of manually matching small patches of cortex within a very limited field of view.

## Validation

Assessing the quality of the registration performance proved challenging. While there is a clear theoretical relationship between the distortion size and the parameters of the acquisition (Jezzard and Balaban, 1995), in practice it is difficult to find a gold standard for the submillimetre accuracy that we are aspiring to. Tools to convert a field maps to estimates of voxel displacement are usually heavily smoothed and cannot account for subject movement in the scanner or field changes between acquisitions. We hence created our own gold standard by acquiring an undistorted FLASH image and manually distorting it based on a field map. This way, the exact distortions were known in order to test if we could find them back with RBR.

A high resolution whole brain multi-echo FLASH image (Haase et al., 1986) was acquired at a 3T Siemens Scanner, TR=95 ms, *α*=20°, bandwidth=170 Hz/px, [0.75 mm]^3^. GRAPPA was used for three-fold in-plane acceleration. The echo times ranged from 5.88 ms to 78.96 ms with an echo spacing of 8.12 ms. The average of the last seven echoes was used, as the first three contained little contrast. This was accompanied by a whole brain MPRAGE acquisition that was used for the FreeSurfer cortical reconstruction, TR/TE/TI/*α* = 2300 ms/3.15 ms/1100 ms/8°, [0.8 mm]^3^. A field map was acquired to realistically distort the FLASH image. The resolution was [3.5 mm X 3.5 mm X 2.0 mm], TR/TE1/TE2/*α* = 1020 ms/10 ms/12.46 ms/90°, bandwidth=260 Hz/Px.

The cortical reconstruction from FreeSurfer’s recon-all (Dale et al., 1999) was coregistered to the FLASH image using a 6 DoF linear registration with a custom MATLAB implementation of BBR. In order to distort the cortical surface, a voxel displacement map (VDM) was computed using SPM field map tools (Andersson et al., 2001). In order to taper unrealistically large displacements, and to move non-displaced vertices away from zero, the VDM was first transformed by a cubic root, after which it was applied in the anterior-posterior direction to the boundaries. Vertices were on average displaced by 2.56 mm (3.4 voxels), which is considerably more than could be tolerated for a laminar specific experiment. The distribution of displacement values was bimodally distributed away from zero with the specific goal to let RBR find the displacement and yield a sharp unimodal distribution around zero, as close to a delta distribution as possible.

Additionally, RBR was tested on 12 brain scans obtained from a 7T scanner. We used 3D EPI (Poser et al., 2010), [0.93 mm]^3^, TR/TE/*α* = 2768 ms/20 ms/14°, bandwidth=1167 Hz/pixel, phase encoding direction: A -> P, matrix size = 204^2^, effective echo spacing = 0.25 ms. The boundaries were created by FreeSurfer on a whole-brain MP2RAGE (1.03 mm^3^, TR/TE/TI1/TI2 = 5000 ms/1.89 ms/900 ms/3200 ms) (Marques et al., 2010). BBR was performed by `bbregister` with a full affine transformation (12 DoFs) on the mean of the functional images and subsequently, the boundaries were imported to MATLAB. Overlaying the registered boundaries on top of the EPI image showed clear local geometrical distortions in the phase encoding direction, related to field inhomogeneity (Jezzard and Balaban, 1995). These distortions were of the order of several millimeters within a single volume. We corrected this with RBR and investigated its performance.

It must be noted that it is challenging to find an objective metric by which the quality of the registration could be quantified. The true distorted position is unknown and methods to approximate do not have the desired submillimetre specificity that is required. Moreover, if such metric existed, it could itself be used in the optimisation procedure. An alternative, however, is an investigation of RBR’s performance on the true volume compared to the performance on the volume with different levels of added noise. Assuming that the algorithm has the best performance on the original data, the displacement with respect to the no-noise condition is expected to increase when more noise is added. Note that this does not make a statement about the correctness of the result. To investigate the RBR’s performance in the presence of noise, we added twelve different levels of white noise to the data, applied RBR and compared the displacement to the unsalted version. For this, we used the Average Absolute Distance (AAD) (Greve and Fischl, 2009), defined to be the average distance that the cortical surface moves between the two sets of registrations.

## Results

We employed RBR on a constructed gold standard where the exact displacement was known. By taking an undistorted FLASH image and realistically distorting the cortical surface by means of a field map, we could test if we could retrieve the initial position of the boundaries. In Fig 2, we present a histogram of the displaced boundaries and the registered boundaries, both with respect to the true position. The registered boundaries clearly show a sharp distribution centred around the origin (*µ* = 0.027 mm). The FWHM of the distribution is 0.49 mm, showing that RBR provides accurate submillimetre registration. Additionally, Fig. 3 shows a cross section (middle slice of the volume) of the registration.

**Figure 2:**
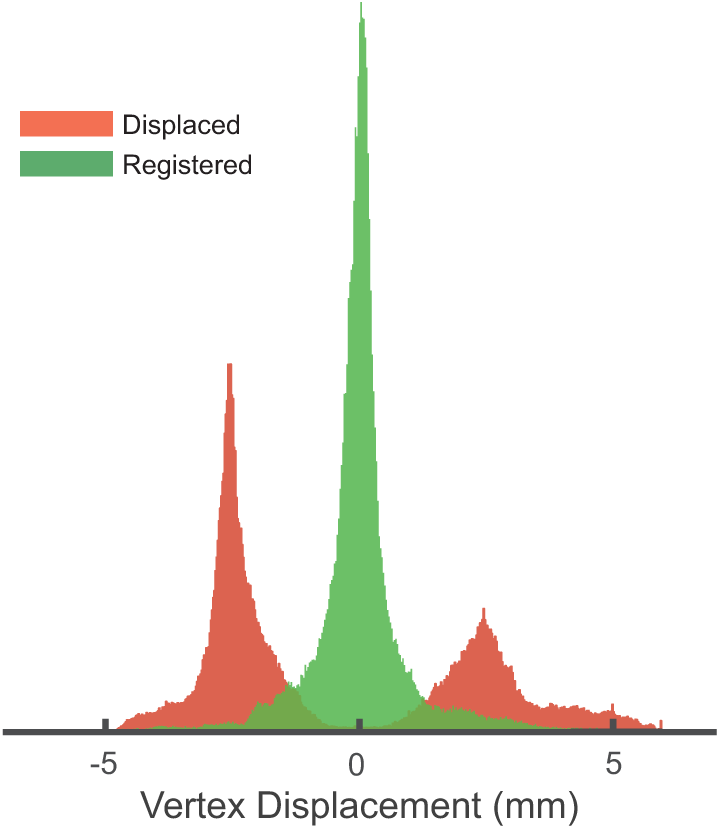
Histogram of the displacement of all 228,208 vertices within a single brain mesh. The red histogram shows the displacement after fieldmap based distortion with respect to a gold standard. The green shows the displacement after applying RBR with respect to the same standard. After registration, the histogram clearly shows a sharp zero-centered (no bias) distribution.

**Figure 3:**
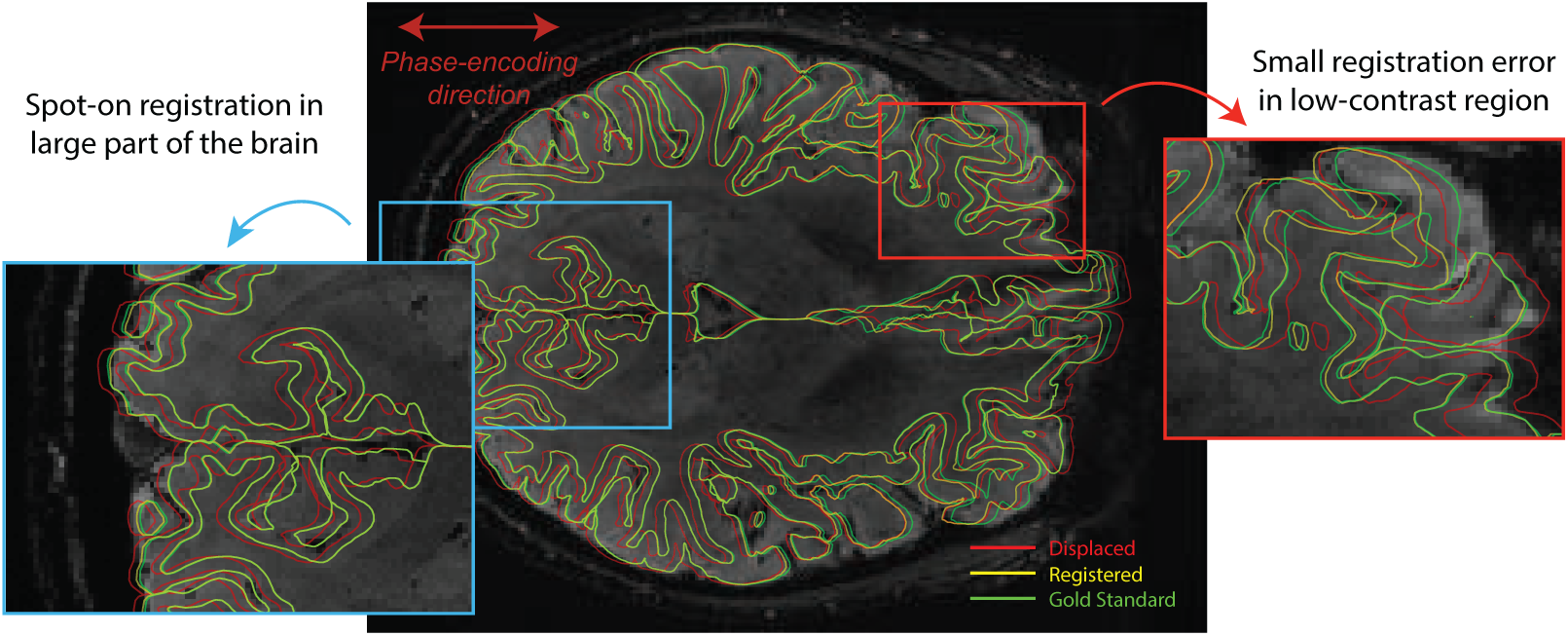
The mid slice of the registration performed on a manually distorted FLASH image (red boundaries). The registered surface (yellow) overlaps for the larger part of the brain almost perfectly with the gold standard (green). If the specificity is set to high values, there is a risk that errors start to appear in some low contrast regions. This largely depends on the balance between false positives and false negatives in terms of corrected regions.

In most of the slice (and the volume), the registration accuracy is well within the submillimetre regime. However, especially in low contrast areas the algorithm may show some small inaccuracies. This mainly proliferates when there is also another gradient in the image (e.g. the pial surface) on which the algorithm starts to fix the boundaries. This is largely related to the fine line between obtaining sufficient specificity and overfitting the data.

We performed RBR on a (resting state) dataset intended for laminar analysis consisting of 11 subjects. The boundaries after a 12 DoF `bbregister` were recursively registered to the mean EPI images (0.93 mm isotropic resolution) and this yielded an updated cortical surface. The new surface followed the grey matter boundary in the volume visibly better than the unregistered one. In Fig. 4 we present a single slice with both sets of boundaries overlaid on top of them, illustrating the improvement. We here present the data for a representative subject, and identical images for all other subjects are presented in the Supplemental Materials.

**Figure 4:**
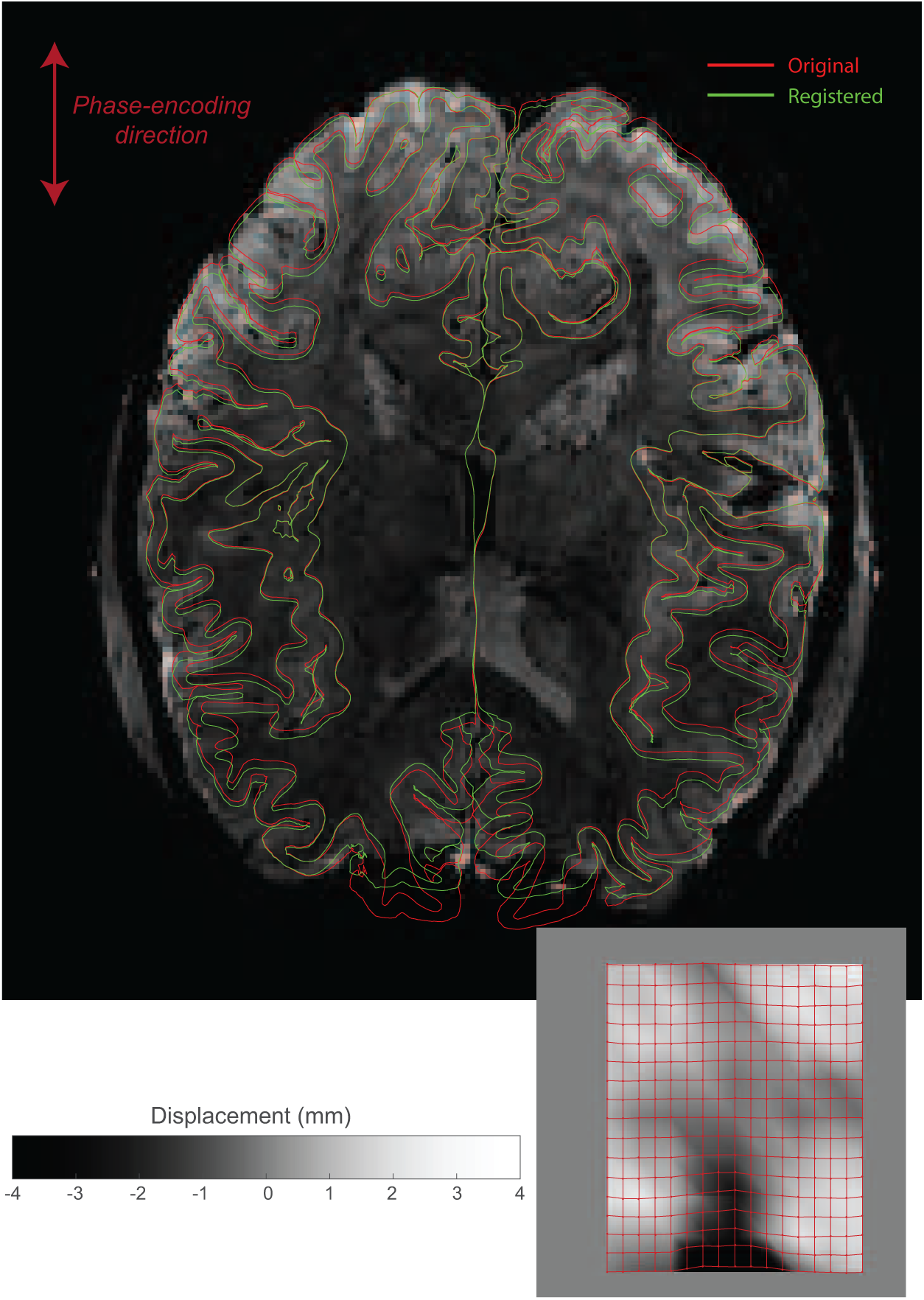
Distortion correction of 7T 3D EPI data, obtained with 0.93 mm isotropic resolution. In red, the original brain surfaces are shown after a 12 DoF registration performed by bbregister. The mesh in green is the updated mesh by means of the first stage RBR, 2 DoF (scale and translation in the PE direction). The green boundaries follow the white matter boundary much better. The voxel displacement map (lower right) shows displacements on the order of several millimetres. The control point lattice that was used to displace the boundaries are overlaid onto the displacement map. Similar images for all subjects can be found in the supplementary materials. For even better inspection, movie files for all subjects are included in the online supplemental materials.

Additionally, Fig 5 shows the Absolute Average Displacement (AAD) of the registration on salted images with respect to the registration on the no-noise volume. Even for highly noisy images, the registration improves somewhat with respect to the undisplaced boundaries. In the absence of a gold standard model of the distortions, the monotonic decrease of the AAD indicates that RBR converges to an optimum as a function of data quality.

**Figure 5:**
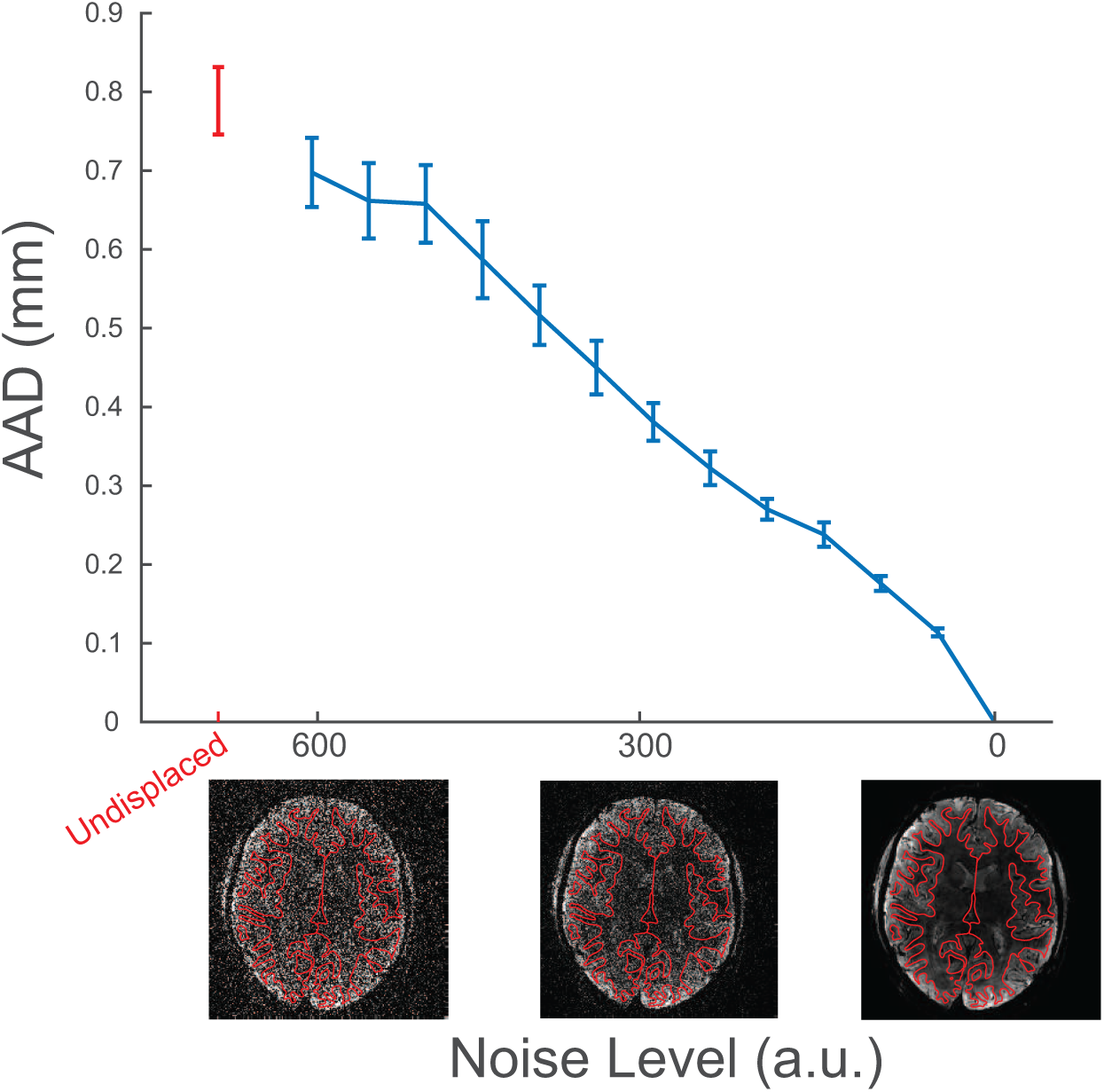
The effect of adding noise on the computed average absolute displacement with respect to a no-noise registration. In the absence of knowledge about the true distortions, the monotonic decrease of the AAD indicates that RBR converges to an optimum as a function of data quality.

**Figure 6:**
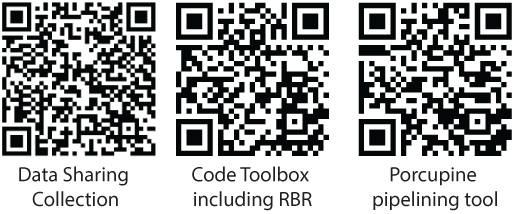
The links to, respectively, the source code of the algorithms, the analysis scripts for creating all figures in this paper, and the pipeline tool to easily incorporate it in one’s custom analysis.

**Figure 7:**
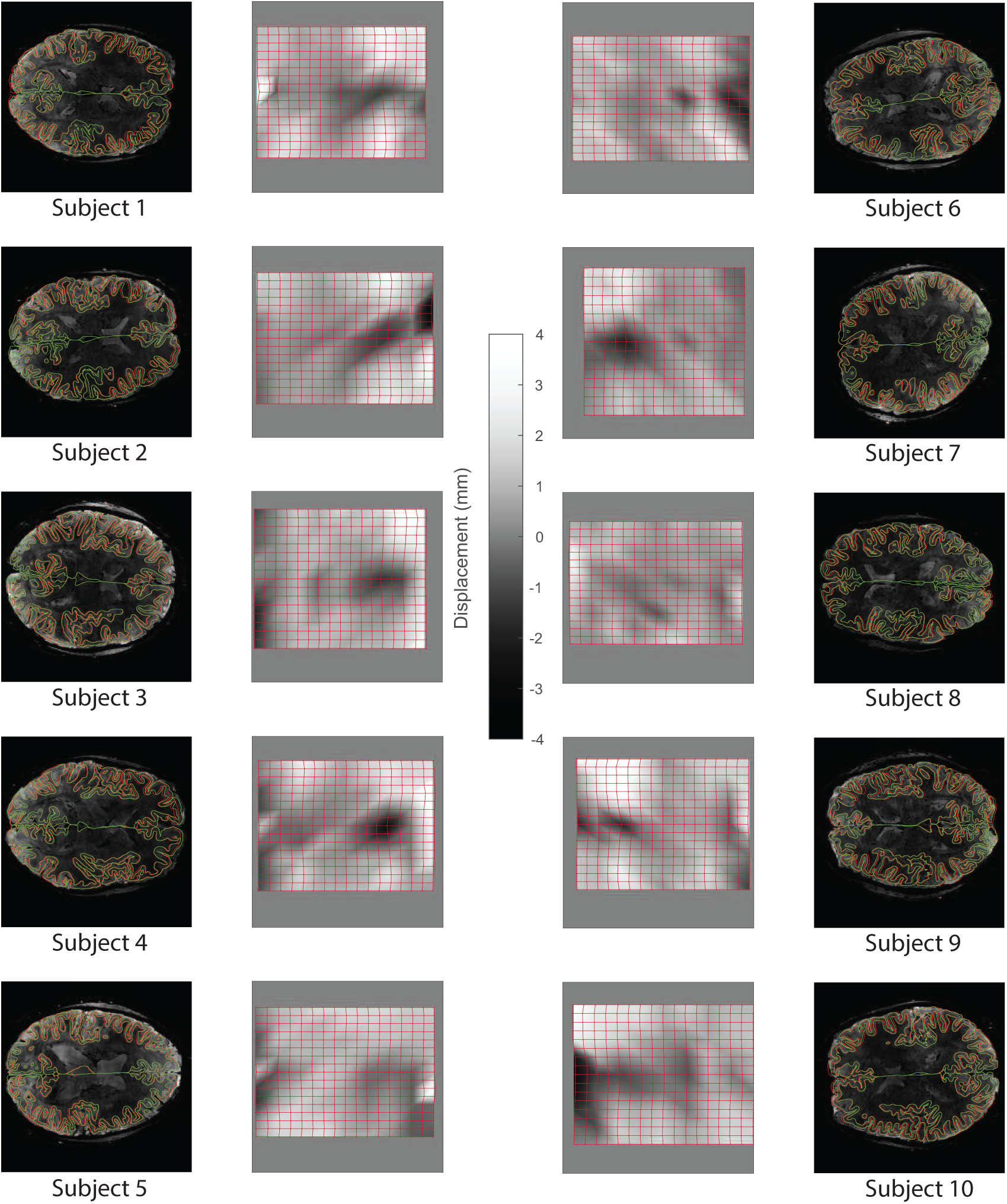
Distortion correction of 7T 3D EPI data, obtained with 0.93 mm isotropic resolution. In red, the original brain surfaces are shown after a 12 DoF registration performed by bbregister. The mesh in green is the updated mesh by means of the first stage RBR, 2 DoF (scale and translation in the PE direction). The green boundaries follow the white matter boundary much better. For better inspection, the online supplemental materials include movie files that scroll through the volume. The voxel displacement map (lower right) shows displacements on the order of several millimetres. The control point lattice that was used to displace the boundaries are overlaid onto the displacement map.

## Data Availability

All source code for the registration algorithm is freely available under the GPL 3.0 license at https://github.com/TimVanMourik/OpenFmriAnalysis. The respective modules are also available in Porcupine https://timvanmourik.github.io/Porcupine, a visual pipeline tool that automatically creates custom analysis scripts (van Mourik et al., 2017). All code to generate the images in this paper are available at the Donders Repository https://data.donders.ru.nl/collections/shared/di.dccn.DSC_3015016.05_558/id/27015532?0. This also includes additional movie files that scroll through the volumes, which is highly relevant to fully investigate whole brain registration performance.

## Discussion and conclusions

In recent years, laminar specific fMRI analysis has come in reach for human in vivo experiments. Where standard fMRI analyses are done on a routine basis, this is hardly the case for layer specific analysis, not least because OF the large amount of manual work that is involved. Aligning volumes to adjust for subject motion and non-linear distortion is one of the most challenging parts of the pipeline. With Recursive Boundary Registration (RBR) we here propose a solution to this problem by means of a recursive application of Boundary Based Registration. By using a control point lattice that forms the basis of deforming the mesh, the topology of the surface is preserved. This way, the mesh can easily be fed into next steps in making a cortical layering (Level Set Method) where it is an absolute necessity that the mesh is not broken as a result of the transformations. Due to the large number of degrees of freedom that is required to non-linearly deform a volume, we built in several methods to increase robustness. Nonetheless, as a mathematical certainty, increasing the sensitivity will increase the false positive rate, i.e. the risk of overfitting. It is up to the user to find the desired balance between them in the settings that we provide to control this balance.

To ensure the functionality of this method, several conditions have to be met. A good reconstruction of the cortical surface is essential, as any inaccuracies will directly affect the contrast estimation and hence the registration optimisation. As the algorithm is based on edge detection, the volume to which it is registered needs to have sufficient contrast between white and grey matter. Furthermore, RBR is recommended to be initialised by a linear BBR registration. While RBR excels at subtle non-linear deformations, large displacement at any level may not be found in a rough multidimensional landscape through which the gradient descent method has to find it way to a minimum. In general, a gradient descent method may be difficult to monitor as it is susceptible to local minima trapping. A more robust algorithm may be a good improvement to RBR (e.g. using bounded gradient descents, or Monte Carlo sampling). Additional points of improvement could be the way in which the control point lattice is applied. While the subdivision into tetrahedra guarantees continuity across segments, it has a non-symmetrical orientation. The resulting displacement hence may show a residual bias as a result of the division. A smoother curve fitting technique to replace the current tetrehedral solution may be a valuable improvement, for example a diffeomorph version as implemented in ANTS (Avants et al., 2011).

This algorithm in its current implementation focuses only on the *gradient* around the gray and the white matter, without looking at the intensity value. One could imagine this to be problematic when the RBR is applied to a volume where a similar gradient of the same direction is present, for example the grey matter to CSF gradient. The algorithm may easily get ‘confused’ about which boundary it needs to converge on. In 
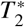
-weighted images in functional studies, this is not usually a problem as the grey matter has a higher intensity value than both CSF and white matter. However, in standard anatomical scans with *T*_1_ contrast, this is not the case. As a result, RBR may fail due to the great similarity of contrast along both sides of the grey matter. An obvious extension for future use could be to use more prior knowledge about the intensity (as apposed to merely the gradient), in order to inform the algorithm which boundary it should use. With the open source and modular way the code is written, the cost function, optimisation algorithm or deformation algorithm could easily be replaced with improved versions, to accommodate ongoing development.

A common use of non-linear registration is template matching, for example in an anatomical-to-MNI normalisation. RBR has little potential for this type of usage as it is based on finding fine within-subject similarities in a volume, rather than the coarser type of between-subject similarities. More probable potential use cases may be could in the matching of subcortical brain structures that are described with a three-dimensional mesh. Similarly, there is a wide range of more deformable parts of the body (e.g. liver, heart, stomach, etc.) that might benefit from solutions like RBR.

We have shown that RBR automates the process of non-linear distortion correction in order to accommodate an extremely high specificity. This is a next step in automated processing of laminar analysis and more routine layer research. We have shown this yields submillimetre accuracy of registration. Finally, we concur with Saad (Saad et al., 2009) and Greve (Greve and Fischl, 2009) in their conclusion to encourage visual inspection of the registration before reliance is placed on its accuracy.

## Supporting Information

These are single slice images of the registration performance for all other 10 subjects. More detailed information can be found in the online supplementary materials.

## Acknowledgements

The authors would like to thank Irati Markuerkiaga for helping with collecting (and recollecting) the FLASH data set. Tim van Mourik acknowledges support by the Spinoza grant [SPI 40-118].

